# Efficient gene disruption with CRISPR-Cas3 in human T cells

**DOI:** 10.1101/2025.08.06.668737

**Authors:** Tomoaki Fujii, Yukimi Sakoda, Kazuto Yoshimi, Kohei Takeshita, Kazumasa Yokoyama, Shoji Watanabe, Koji Tamada, Tomoji Mashimo

## Abstract

The CRISPR-Cas9 system has been widely adopted as a genome editing tool due to its high efficiency and versatility, contributing to the development of various therapeutic strategies. However, its clinical application remains limited by safety concerns, including off-target effects and large-scale chromosomal rearrangements such as translocations and inversions. Recently, the CRISPR-Cas3 system, a Class 1 CRISPR effector complex with unidirectional DNA degradation activity, has gained attention as a potential alternative, offering reduced off-target activity. In this study, we applied the CRISPR-Cas3 system to human T cells and successfully disrupted two clinically relevant genes, T cell receptor alpha constant (TRAC) and beta-2 microglobulin (B2M). These gene deletions were associated with a reduction in both graft-versus-host disease (GVHD) risk and host immune rejection. Importantly, no off-target mutations were detected in CRISPR-Cas3-edited cells, in contrast to the off-target effects observed with CRISPR-Cas9. Furthermore, CAR-T cells generated by deleting *TRAC* or *B2M* using CRISPR-Cas3 maintained their antigen-specific cytotoxicity against tumor cells, while exhibiting reduced alloreactivity. These results suggest that CRISPR-Cas3 provides a safer and promising platform for genome editing in T cell engineering, with potential applications in the development of next-generation allogeneic T cell therapies.

## Introduction

Genome editing technologies have transformed biomedical research and therapeutic development. Among these, the CRISPR (clustered regularly interspaced short palindromic repeats)-associated protein systems, originally discovered as part of the prokaryotic adaptive immune system [1], have become powerful tools for site-specific genome modification. Owing to their programmability and high efficiency, CRISPR-based approaches—particularly those employing Class 2 single-effector nucleases—have been widely adopted in both research and clinical settings. To date, most genome editing in mammalian cells has utilized Class 2 systems, including Cas9 (Type II) [2], Cas12 (Type V) [3], and Cas13 (Type VI) [4]. Among these, Cas9 has demonstrated robust editing activity in eukaryotic cells and has facilitated the development of novel therapeutic strategies [5, 6]. Nonetheless, Cas9-mediated genome editing carries safety concerns, such as off-target effects and chromosomal rearrangements including translocations and inversions.

In contrast, Class 1 CRISPR systems utilize multi-component effector complexes and have been less frequently applied in mammalian genome editing. One such system, CRISPR-Cas3 (Class 1, Type I-E), induces long-range, unidirectional DNA degradation rather than small insertions or deletions [7]. This system operates through a multi-step mechanism: Cas6 processes the CRISPR RNA (crRNA), which is then incorporated into the Cascade surveillance complex, composed of Cas5, Cas6, Cas7, Cas8, and Cas11. Upon target recognition and R-loop formation, the complex recruits Cas3, a helicase-nuclease that degrades DNA processively [8]. A notable feature of CRISPR-Cas3 is its enhanced target specificity, attributed to its longer crRNA recognition sequence (27 nucleotides) compared to that of Cas9 (20 nucleotides). This extended sequence may contribute to reduced off-target activity, making Cas3 an attractive candidate for precise genome editing applications, especially where safety is a critical concern.

One promising application of CRISPR-Cas3 is in the development of allogeneic T cell therapies, also referred to as universal CAR-T cells. While several autologous CAR-T cell therapies have been approved [9], their dependence on patient-derived T cells presents challenges in terms of manufacturing and logistics. Universal CAR-T therapies, derived from healthy donors, offer a more scalable alternative but require genome editing to eliminate endogenous T cell receptor (TCR) and HLA class I molecules to avoid graft-versus-host disease (GVHD) and immune rejection. Although Cas9 has been used to achieve these modifications, safety concerns remain regarding its off-target activity. Despite its potential advantages, the application of CRISPR-Cas3 in human T cells has not been extensively explored. In this study, we investigated the feasibility of using CRISPR-Cas3 to generate allogeneic T cells by targeting TRAC and B2M. By optimizing crRNA design and delivery methods, we achieved efficient gene disruption with no detectable off-target effects. These results highlight the potential of CRISPR-Cas3 as a safer genome editing platform for the development of universal T cell therapies.

## Results

### crRNA screening for CRISPR-Cas3-mediated *TRAC* and *B2M* disruption

Deletion of *TRAC* and *B2M* is a well-established strategy to ablate TCR and major histocompatibility complex class I (MHC-I; HLA-I) surface expression, respectively [10]. To achieve efficient disruption of these genes using CRISPR-Cas3, we performed an *in silico* screen of candidate crRNA sequences based on the following criteria: the presence of a 5′-AAG-3′ protospacer adjacent motif (PAM), a GC content between 30% and 70%, and a target site located 100–500 bp downstream of the target exon. In addition, candidate sequences were filtered to ensure that the full 5′-PAM plus protospacer sequence (allowing up to three mismatches), as well as the 5′-PAM plus 15-nt seed region, showed no significant matches elsewhere in the human genome apart from *TRAC* or *B2M*. The screened crRNAs were co-electroporated into Jurkat cells along with plasmids encoding Cas3, Cascade complex proteins, and each candidate crRNA (Figure 1A). Five days post-electroporation, surface expression of TCR (for *TRAC*) and HLA-I (for *B2M*) was evaluated by flow cytometry (Figure 1B). Among the tested sequences, the most efficient crRNAs mediated a maximum of 20.0% and 11.3% disruption of *TRAC* and *B2M* expression, respectively. Based on these results, we selected three top-performing crRNAs for each gene (*TRAC* #3, #5, #9 and *B2M* #3, #15, #20) for further study.

**Figure 1.**
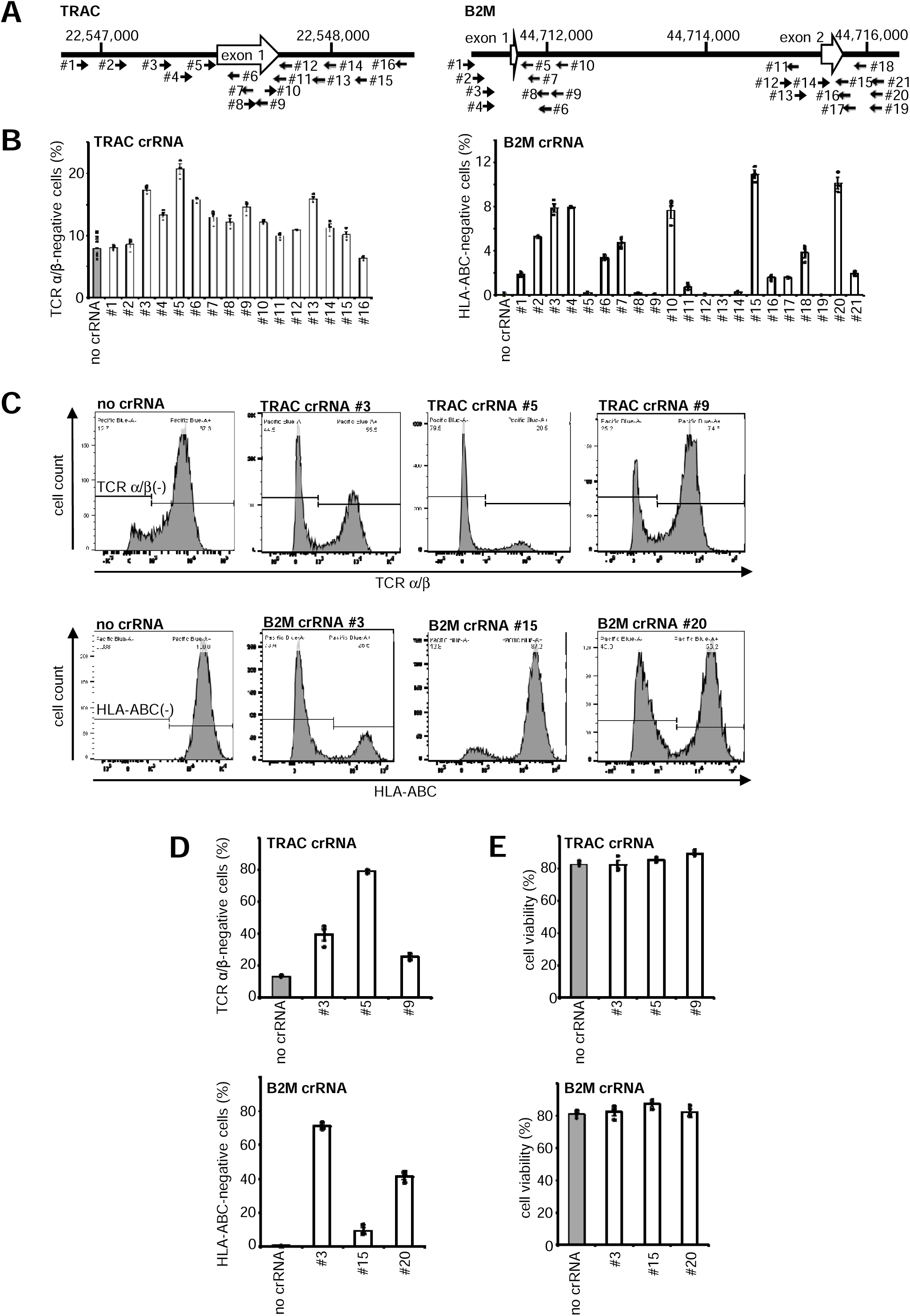
Selection of crRNAs targeting *TRAC* and *B2M* for CRISPR-Cas3–mediated gene disruption in Jurkat cells. (A) Schematic illustration of crRNA target sites on the *TRAC* and *B2M* loci. The selected target sequences are located within exon 1 of *TRAC* and exons 1 and 2 of *B2M*. (B) Quantification of TCRα/β and HLA-ABC disruption efficiency by Cas3-Cascade-crRNA plasmid electroporation, measured by flow cytometry (mean ± SEM, n[≥[3). (C) Representative flow cytometry plots showing TCRα/β and HLA-ABC expression in Jurkat cells following CRISPR-Cas3 editing. (D) Summary of gene disruption efficiency (mean ± SEM, n[=[3). (E) Cell viability following electroporation (mean ± SEM, n[=[3).

Previous studies have demonstrated that delivery of Cas9 as a ribonucleoprotein (RNP) complex enables highly efficient genome editing in human T cells [11–13]. Building on this concept, we generated Cas3-Cascade/crRNA RNP complexes corresponding to the selected crRNA sequences [8, 14]. These RNPs were electroporated into Jurkat cells, and functional gene disruption was evaluated five days later. Disruption efficiencies for *TRAC* were 39.5% (crRNA #3), 79.0% (#5), and 25.4% (#9), and for *B2M* were 71.1% (crRNA #3), 9.4% (#15), and 41.3% (#20) (Figure 1C, D). Importantly, cell viability at day 5 post-electroporation was comparable between Cas3-RNP treated and untransfected cells (Figure 1E, Table S1), indicating that Cas3-Cascade RNP delivery does not compromise cell viability in Jurkat cells. Collectively, these results demonstrate that the CRISPR-Cas3 system enables efficient and specific disruption of *TRAC* and *B2M* genes in human T cells without impairing cell viability.

### No off-target and tumorigenic mutations with CRISPR-Cas3

CRISPR-Cas technologies are widely recognized as powerful tools for gene therapy due to their high editing efficiency; however, concerns about off-target effects remain a major barrier to clinical translation [15–18]. As previously reported, CRISPR-Cas3 possesses a longer target recognition sequence compared to Cas9, which is expected to reduce the risk of off-target activity [7]. To experimentally evaluate the off-target risks associated with Cas3-Cascade RNP delivery, we performed capture sequencing of predicted potential off-target (POT) sites in the human genome for *TRAC* crRNAs (#3, #5, #9) and *B2M* crRNAs (#3, #15, #20) ( The detailed information is not disclosed. ). POT sites were identified based on a previously published scoring algorithm [7]. Off-target scores were calculated by subtracting the number of split reads detected in negative control cells (electroporated with Cas3-Cascade RNPs lacking crRNA) from those in cells transfected with crRNA-containing RNPs. Elevated scores were observed exclusively at the intended on-target regions, while no significant signals were detected at any other POT sites across the genome (Figure 2A). These findings indicate that Cas3-mediated genome editing did not result in detectable off-target mutations at any of the examined loci.

**Figure 2.**
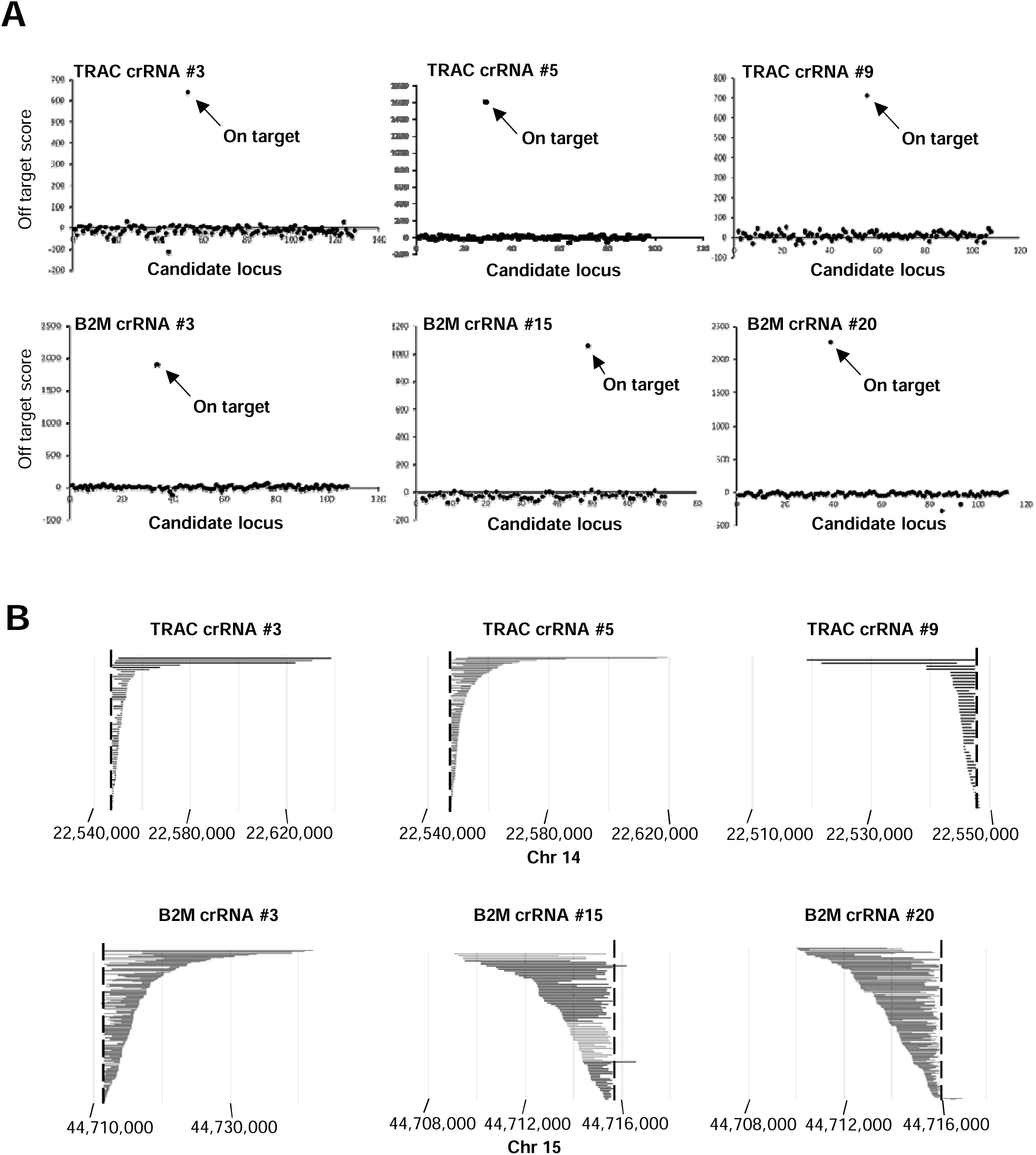
CRISPR-Cas3 induces no detectable off-target or tumorigenesis-related mutations. (A) Off-target analysis in Jurkat cells transfected with Cas3-Cascade RNPs. Deep sequencing was performed using custom microarray-based capture of on- and off-target regions. No off-target deletions were detected at predicted potential off-target (POT) sites. (B) Deletion patterns induced by Cas3-Cascade RNP in Jurkat cells. Each horizontal bar represents a deletion event aligned at the distal cleavage site. The dashed line indicates the position of the crRNA. The deleted genomic regions surrounding *TRAC* and *B2M* did not include known tumor suppressor genes.

Given the processive nature of Cas3-mediated DNA degradation [2, 3], we next assessed the extent of large genomic deletions and their potential impact on neighboring genes, including those potentially involved in tumorigenesis. To this end, we performed capture sequencing across 100-kbp regions flanking both the 5′ and 3′ ends of the *TRAC* and *B2M* loci. The largest observed deletion at the *TRAC* locus extended 66.8 kbp in the 3′ direction (crRNA #5) and 28.3 kbp in the 5′ direction (crRNA #9). At the *B2M* locus, deletions of up to 30.5 kbp (3′ side, crRNA #3) and 5.8 kbp (5′ side, crRNA #15) were observed (Figure 2B and Table S2). To evaluate whether these extended deletions affected neighboring genes, we examined loci adjacent to the target sites. *ABHD4* and *DAD1* are located near the *TRAC* locus, while *PATL2* and *TRIM69* flank the *B2M* locus. In cells transfected with *TRAC* crRNAs, we identified 66 (crRNA #3), 154 (#5), and 49 (#9) large deletion patterns. Among these, partial deletions of the *DAD1* gene were found in 7.5% (5/66) and 5.2% (8/154) of cases with crRNAs #3 and #5, respectively. *ABHD4* was affected in 4.4% (3/66, crRNA #3) and 1.3% (2/154, crRNA #5), while no adjacent gene deletions were detected with crRNA #9. For *B2M* crRNAs, 176 (crRNA #3), 87 (#15), and 174 (#20) large deletion events were identified. Among these, partial deletions of *PATL2* were observed in 12.7% (11/87, crRNA #15) and 6.9% (12/174, crRNA #20), while *TRIM69* was affected in 2.3% (4/176, crRNA #3) (Figure 2B and Table S2). Together, these results demonstrate that CRISPR-Cas3 enables highly specific genome editing in human T cells without detectable off-target effects, and that while rare extended deletions into neighboring genes may occur, no genes with known tumorigenic potential were disrupted. These findings support the potential of CRISPR-Cas3 as a precise and safe genome editing modality for therapeutic applications.

### Efficient genome editing in primary T cells by CRISPR-Cas3 mRNAs

To carry out functional allogenic CAR-T assay, we attempted to delete *TRAC* and *B2M* genes with Cas3-Cascade RNP in human primary T cells. However, its results were poorly reproducible possibly due to poor cell viability after electroporation of RNP (data not shown). Recent advancement of mRNA therapeutic modality motivated us to examine its applicability to Cas3 genome editing. We therefore synthesized Cas3 and Cascade mRNAs with a Cap1 structure and uridine replaced by N1-methylpseudouridine (N1-me-Ψ). The crRNA containing 29 nt 5’- and 3’-repeat and 32 nucleotide spacer sequences were synthesized with 2’-O-methyl phosphorothioate linkage modification at the 5’- and 3’-terminal 3 nucleotides. We transfected these modified mRNA and crRNA (*TRAC* crRNA #5 and *B2M* crRNA #3) into human primary T cells and evaluated functional disruption efficiency with flow cytometry 96 hours after electroporation (Figure S1). We systematically optimized electroporation conditions and finally observed that *TRAC* and *B2M* were eliminated in 34.7% and 45.0% in human primary T cells, respectively (Figure 3A and B). Notably, cell viability was preserved across all tested samples (Figure 3C and Table S3).

**Figure 3.**
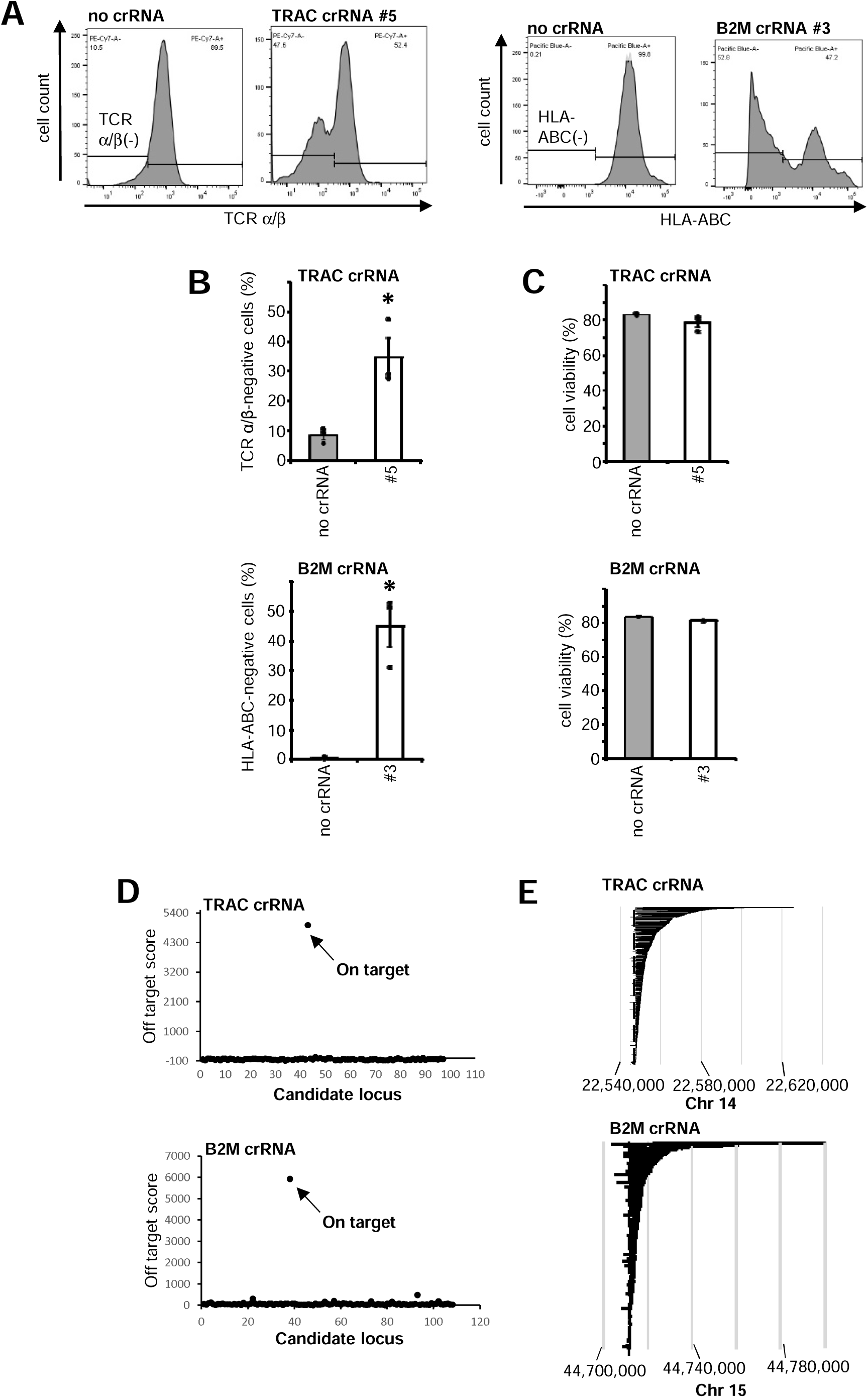
Highly efficient *TRAC* and *B2M* disruption in human primary T cells using CRISPR-Cas3 mRNA without compromising cell viability. (A) Representative flow cytometry plots showing expression of TCRα/β and HLA-ABC in primary T cells electroporated with Cas3-Cascade mRNA and crRNA targeting *TRAC* or *B2M*. (B) Quantification of gene disruption efficiency for *TRAC* and *B2M* (mean ± SEM, n = 3). Statistical significance was determined by Student’s t-test. (C) Cell viability following Cas3-mediated editing (mean ± SEM, n = 3). (D) Off-target analysis by capture-based deep sequencing showed no detectable off-target mutations in edited T cells. (E) On-target deletion analysis confirmed precise and directional genome excision at *TRAC* and *B2M* loci. A deletion event at the distal cleavage site is represented by each horizontal bar. The location of the crRNA is indicated by the dashed line.

We subsequently preformed off- and on-target analysis as described above. The analysis demonstrated no apparent off-target mutations with the CRISPR-Cas3 (Figure 3D). On-target analysis revealed 746 deletion patterns for *TRAC* crRNA #5 and 736 patterns for *B2M* crRNA #3. The maximum deletion sizes reached 77.3 kbp (3′ side) and 10.1 kbp (5′ side) at the *TRAC* locus, and 84.7 kbp (3′ side) and 13.1 kbp (5′ side) at the B2M locus (Figure 3E). Among the *TRAC* deletions, 8.7% (65/746) included partial loss of the neighboring *DAD1* gene, and 0.7% (5/746) affected *ABHD4*. For *B2M*, 7.6% (56/736) of deletions involved *PATL2*, and 2.6% (19/736) involved *TRIM69* (Table S4).

To benchmark these results, we evaluated off-target activities of the CRSIPR-Cas9 system in human T cells. Adjusted indel percentages were calculated by subtracting frequency of insertion and deletion frequencies in control cells introduced with Cas9 alone (without sgRNA) from those in cells transfected with Cas9 and target-specific sgRNAs. The insertion and deletion counts were determined by using Crispresso2 [19]. The adjusted indel values revealed the percentage of off-target mutations in both *TRAC*- and *B2M*-deleted human T cells generated with CRISPR-Cas9 system (Figure S2). Taken together, these results demonstrate that the CRISPR-Cas3 system enables efficient and precise disruption of *TRAC* and *B2M* in human primary T cells without compromising cell viability or inducing detectable off-target effects. In contrast, CRISPR-Cas9 editing at the same loci was associated with measurable off-target activity.

### Reduced alloreactivity and preserved tumor killing in Cas3-engineered CAR-T cells

Peripheral blood mononuclear cells (PBMCs), with *TRAC* or *B2M* disrupted using the Cas3 mRNA and modified crRNA method described above, were cultured for 5 days. T cells were then magnetically enriched based on *TRAC* or *B2M* surface expression to isolate gene-disrupted (negative) and non-disrupted (positive) populations (Figure 4A). These enriched T cells were then transduced with a retroviral vector encoding an anti-GM2 CAR and further expanded for 4 days. Flow cytometric analysis confirmed that both *TRAC*- and *B2M*-negative CAR-T cells expressed CAR on the cell surface at levels comparable to their respective positive counterparts, while maintaining suppression of *TRAC* or *B2M* expression (Figure 4B).

**Figure 4.**
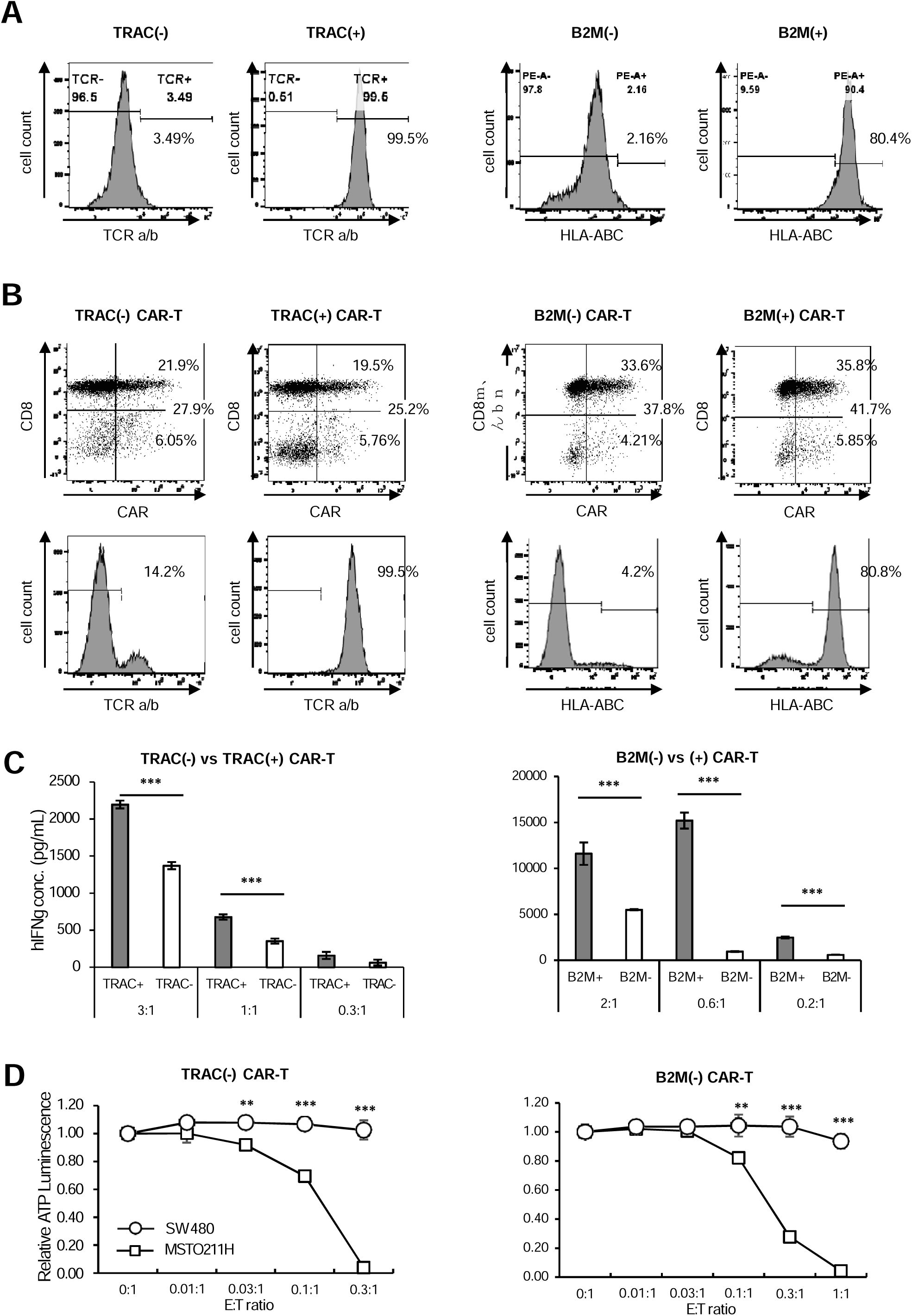
CRISPR-Cas3–engineered CAR-T cells retain cytotoxic function and exhibit reduced alloreactivity. (A) Flow cytometric analysis of TCRα/β and HLA-ABC expression in *TRAC*- or *B2M*-disrupted versus non-disrupted T cells after magnetic cell sorting. (B) CAR expression and residual TCRα/β or HLA-ABC expression in *TRAC*- or *B2M*-disrupted versus non-disrupted CAR-T cells assessed by flow cytometry. (C) Mixed lymphocyte reaction (MLR) assays evaluating alloreactivity of *TRAC*-disrupted CAR-T cells co-cultured with irradiated allogeneic PBMCs, or *B2M*-disrupted CAR-T cells co-cultured with allogeneic T cells. Data represent mean ± SD from triplicate wells (n = 3). (D) Cytotoxic activity of *TRAC*- or *B2M*-disrupted CAR-T cells against GM2-positive or GM2-negative tumor cells. Data represent mean ± SD from triplicate wells (n = 3). Statistical significance was determined using unpaired two-tailed Student’s t-test: *p < 0.05, **p < 0.01, ***p < 0.001.

To assess alloreactive potential, we performed mixed lymphocyte reaction (MLR) assays using *TRAC*- or *B2M*-positive and -negative CAR-T cells as both responders and stimulators. *TRAC*-negative CAR-T cells produced significantly lower levels of IFN-γ in co-culture with irradiated allogeneic PBMCs compared to *TRAC*-positive controls (Figure 4C), indicating reduced alloreactivity as responder cells. Similarly, *B2M*-negative CAR-T cells induced lower IFN-γ secretion from co-cultured allogeneic T cells, demonstrating that loss of HLA class I molecules attenuates stimulation of host T cell responses when used as stimulator cells (Figure 4C). Importantly, the anti-tumor cytotoxic activity of *TRAC*- or *B2M*-disrupted CAR-T cells, assessed via *in vitro* tumor cell killing assays, remained comparable to that of non-edited CAR-T cells (Figure 4D). Collectively, these results demonstrate that CRISPR-Cas3-mediated disruption of *TRAC* and *B2M* reduces alloreactive immune responses while preserving anti-tumor function, underscoring the utility of Cas3-based editing for universal CAR-T cell therapies.

## Discussion

CRISPR-Cas systems have emerged as powerful tools for genome engineering and are being explored for the clinical manufacturing of allogeneic CAR-T cells. Multiple studies have demonstrated successful and efficient disruption of immunologically relevant genes using the CRISPR-Cas9 system [20–22]. While Cas9-based editing is highly efficient, off-target effects have been widely reported and remain a major concern in therapeutic applications due to the risk of unintended consequences, including potential oncogenesis [18]. Thus, a genome editing platform with both high precision and minimal off-target activity would be highly advantageous for clinical translation. We have previously reported that CRISPR-Cas3 enables long-range genome deletions with high specificity in human cell lines without detectable off-target mutations [7]. In the present study, we successfully applied the CRISPR-Cas3 system to delete two clinically relevant genes, *TRAC* and *B2M*, in human T cells, demonstrating its feasibility as a genome editing tool for generating allogeneic CAR-T cells.

We first designed highly specific crRNAs for *TRAC* and *B2M* using *in silico* screening based on defined criteria, and confirmed their functionality in Jurkat cells using plasmid-based delivery. The most effective crRNAs were selected for further use in Cas3-Cascade RNPs, which enabled highly efficient gene disruption in Jurkat cells. Importantly, capture sequencing confirmed the absence of off-target mutations. These steps can be readily adapted to other gene targets. The on-target deletions extended up to 30.5 kb downstream and 5.8 kb upstream of *B2M*, and up to 66.8 kb downstream and 28.3 kb upstream of *TRAC*, involving regions that contain *PATL2*, *TRIM69*, *DAD1*, and *ABHD4*. *PATL2* is involved in oocyte maturation [23], *TRIM69* is an E3 ubiquitin ligase downregulated in cataract lenses [24], *DAD1* is part of the oligosaccharyltransferase complex and upregulated in certain tumors [25], and *ABHD4* regulates N-acylphospholipids in the central nervous system [26]. None of these genes are tumor suppressors, supporting the safety of our crRNA selections in the context of therapeutic editing.

In human primary T cells, the use of chemically modified mRNA and crRNA enabled reproducible CRISPR-Cas3–mediated gene disruption (Figure 3). Although RNP-based editing using Cas9 is known for its rapid action and reduced off-target effects [27], our attempts to deliver Cas3-Cascade as RNPs in primary T cells were unsuccessful, likely due to incompatibility between the electroporation buffer and RNP stability. We observed aggregation of Cas3-Cascade RNPs, similar to what has been reported with Cas9 RNPs [28]. Optimization of delivery conditions tailored to Cas3 RNP stability will be an important area for future research.

Using the chemically modified mRNA/crRNA approach, we demonstrated that CRISPR-Cas3 induces no detectable off-target mutations in human primary T cells, in contrast to the off-target effects observed with the CRISPR-Cas9 system. These findings provide strong evidence supporting the superior genomic specificity and safety profile of CRISPR-Cas3 for therapeutic genome editing. We successfully disrupted *TRAC* and *B2M* using this mRNA-based platform, achieving efficient gene knockout in primary T cells. Although editing efficiency was slightly lower than that observed in RNP-transfected Jurkat cells, gene-disrupted T cells could be readily purified via magnetic sorting (Figure 4). These edited cells were further engineered into CAR-T cells, and functional assessments demonstrated that *B2M* disruption attenuated host-versus-graft responses, while *TRAC* disruption reduced graft-versus-host alloreactivity—both critical for allogeneic T cell therapy. Importantly, the cytotoxic function of the CAR-T cells was preserved despite dual gene disruption, as shown by *in vitro* tumor killing assays.

Together, our results indicate that CRISPR-Cas3–mediated genome editing of *TRAC* and *B2M* offers a viable strategy for generating functionally competent, allogeneic CAR-T cells with reduced immunogenicity and high genomic precision. This system represents a valuable addition to the genome editing toolkit for allogeneic CAR-T cell production and other therapeutic applications requiring efficient and safe gene disruption.

## Methods

### Plasmids

The plasmids pPB-CAG-hCas3 (Addgene #134920) and pCAG-All-in-one-hCascade (Addgene #134919) were used to express human Cas3 and the Cascade complex, respectively. crRNA expression vectors were generated by cloning 32-bp double-stranded oligonucleotides into the target site of the backbone vector pBS-U6-crRNA-empty (Addgene #134921).

### Cas3-Cascade RNP production

The Cas3 protein, fused to an N-terminal His-tag and a nuclear localization signal (NLS) at either the N- or C-terminus, was produced using the Bac-to-Bac baculovirus expression system (Thermo Fisher Scientific) in *Spodoptera frugiperda* (Sf9) cells, as previously described [8]. Purification was performed using nickel affinity chromatography followed by size exclusion chromatography.

For preparation of the Cascade–crRNA ribonucleoprotein (RNP) complex, the Cascade operon (*cas8–cas11–cas7–cas5–cas6*) was engineered to separate between *cas7* and *cas5*, and an NLS sequence was appended to the C-terminus of each subunit. The respective gene fragments were cloned into the dual multiple cloning sites of the pRSFDuet-1 vector. The pre-crRNA construct (repeat–spacer–repeat) was inserted into the first multiple cloning site of pACYCDuet-1. To co-express the complete Cascade–crRNA complex, pRSFDuet-1 (Cascade operon), pACYCDuet-1 (pre-crRNA), and pCDFDuet-1 harboring His-tagged *cas11* were co-transformed into *E. coli* JM109 (DE3). RNP complexes were purified via nickel affinity and size exclusion chromatography.

### Synthesis of crRNA, sgRNA, and mRNAs for Cas3-Cascade and Cas9

Chemically modified mRNAs for Cas3, Cascade subunits, and Cas9 were synthesized with a Cap1 structure and incorporation of N1-methylpseudouridine (N1-me-Ψ) in place of uridine (Elixirgen Scientific, Inc.) (Table S5). Chemically modified crRNAs for Cas3 and sgRNAs for Cas9 were synthesized with 2′-O-methyl phosphorothioate linkages at the three terminal nucleotides on both the 5′ and 3′ ends (Integrated DNA Technologies, Inc.).

### Electroporation for Jurkat cells

Jurkat cells were electroporated using a 4D-Nucleofector X Unit (Lonza, V4XC-1032) according to the manufacturer’s protocol. Cells were harvested by centrifugation (90 × g, 10 min, room temperature) and resuspended at a density of 2 × 10[ cells in 20 μL of SE Cell Line Nucleofector X solution (Lonza). For plasmid-based delivery, cells were mixed with 500 ng each of pPB-CAG-hCas3, pCAG-All-in-one-hCascade, and crRNA expression plasmids. Electroporation was performed using the CL120 program.

For RNP-based delivery, cells were mixed with 116 pmol each of purified Cas3 protein and Cascade–crRNA complex, and electroporated using the CM137 program. Following electroporation, cells were incubated at room temperature for 30 minutes and then transferred to culture medium at 37 °C in a 5% CO[ incubator. After 120 hours, cells were collected for genomic DNA extraction and flow cytometric analysis.

### Electroporation for primary T cells

Peripheral blood mononuclear cells (PBMCs) were obtained from Charles River Laboratories International, Inc. Cells were thawed and activated for 72 hours on plates pre-coated with anti-human CD3 antibody (OKT3, eBioscience #16-0037-85) and Retronectin (Takara Bio #T100B). T cells were cultured in OpTmizer™ CTS™ T-Cell Expansion SFM (Gibco #A10221-01) supplemented with 2% CTS™ Immune Cell SR (Gibco #A25961-02), 2 mM L-glutamine (Gibco #25030-081), 1% penicillin-streptomycin (Wako #168-23191), 2.5 μg/mL fungizone (Bristol Myers Squibb #4987279112053), and 200 IU/mL recombinant human IL-2 (Peprotech #200-02).

After activation, cells were resuspended at 2.5 × 10[ cells in 25 μL of MaxCyte electroporation buffer (MaxCyte, Inc.). For CRISPR-Cas3 delivery, cells were mixed with 1.0 μg of mRNA encoding each of Cas3, Cas5, Cas6, Cas7, Cas8, and Cas11, along with 1.6 μg of chemically modified crRNA, and electroporated using the MaxCyte GTx instrument with the Expanded T Cell 4 program. For CRISPR-Cas9 delivery, 2.5 μg of Cas9 mRNA and 5 μg of sgRNA were used under identical conditions. Following electroporation, cells were incubated at 37 °C in 5% CO[ for 30 minutes in Retronectin-coated plates. After 24 hours, cells were transferred to fresh Retronectin-coated plates and cultured for an additional 48 hours. Cells were harvested at 72 hours post-electroporation for flow cytometric analysis.

### Generation of anti-GM2 CAR-T cells

Anti-GM2 CAR-T constructs were generated as previously described [29]. Briefly, a humanized anti-GM2 single-chain variable fragment (scFv) was fused to the transmembrane domain of human CD8α and the intracellular signaling domains of CD28, 4-1BB, and CD3ζ. The construct was cloned into a retroviral vector for transduction into human T cells, as described in the same report. CAR expression was assessed by flow cytometry using an anti-idiotype monoclonal antibody specific to the GM2 CAR.

### Flow cytometry

The following monoclonal antibodies (mAbs) were used for analysis: Brilliant Violet 421 anti-human HLA-ABC (BD Biosciences #565332), PE/Cyanine7 anti-human TCR α/β (BioLegend #306719), and Brilliant Violet 421 anti-human TCR α/β (BioLegend #306722). CAR expression was detected using APC anti-human CD8 (BioLegend #300912) and biotinylated anti-idiotype antibody against GM2 CAR (Cell Engineering Corporation), followed by Brilliant Violet 421-conjugated streptavidin (BioLegend #402226). Data were acquired on a BD FACSCanto II or CytoFLEX S (Beckman Coulter) and analyzed using FlowJo software (FlowJo LLC).

### Magnetic sorting

After 5 days of culture, *TRAC*- or *B2M*-disrupted cells were enriched for TCR α/β- or HLA class I-negative populations using magnetic-activated cell sorting (MACS). Briefly, cells were stained with PE-conjugated anti-TCR α/β or PE-conjugated anti-HLA-ABC antibody (BioLegend) for 30 minutes at 4 °C. After washing with MACS buffer (PBS supplemented with 1% heat-inactivated fetal calf serum), cells were incubated with anti-PE MicroBeads (Miltenyi Biotec) for 15 minutes at 4 °C. Cells were then washed, resuspended in MACS buffer, and subjected to positive or negative selection using LS or LD MACS columns (Miltenyi Biotec) according to the manufacturer’s protocol.

### Mixed Lymphocyte Reaction assay

TRAC-positive or -negative CAR-T cells were used as responder (effector) cells and co-cultured with irradiated (30[Gy) peripheral blood mononuclear cells (PBMCs) from a different healthy donor at a density of 2[×[10[ cells per well in a 96-well round-bottom plate, at the indicated effector-to-target (E:T) ratios. For B2M-based stimulation, B2M-positive or -negative CAR-T cells were irradiated (30[Gy) and plated at a density of 3[×[10[ cells per well as stimulator cells, co-cultured with allogeneic T cells isolated using the Pan T Cell Isolation Kit (Miltenyi Biotec), also at the indicated E:T ratios. After 48 hours of incubation, interferon-γ (IFN-γ) levels in the culture supernatant were quantified using the ELISA MAX™ Deluxe Set Human IFN-γ kit (BioLegend), according to the manufacturer’s instructions.

### Cell killing assay

GM2-positive (MSTO-211H) or GM2-negative (SW480) tumor cells were seeded at a density of 5[×[10[ cells per well in 96-well plates and co-cultured with CAR-T cells—with or without CRISPR-Cas3-mediated disruption of TRAC and B2M genes—at the indicated effector-to-target ratios. After 72 hours, tumor cell viability was assessed using the CellTiter-Glo® Luminescent Cell Viability Assay kit (Promega, G7571). Luminescence signals were measured and normalized to wells containing tumor cells alone (negative control) to calculate relative viability.

### Deep sequencing for on- and off-target analysis

Potential off-target (POT) sites for each crRNA used in the CRISPR-Cas3 system were identified in the human genome assembly (GRCh38/hg38), based on previously reported algorithms with slight modifications[7]. Briefly, POT regions were defined as genomic loci containing 16–20 nt consecutive matches, including the PAM sequence, relative to each target site, or as regions tolerating up to 7 mismatches with the full 32-nt target sequence. For CRISPR-Cas9, POT sites were defined as regions allowing up to 3 mismatches. To evaluate on-target editing, probes were designed to cover 100 kbp upstream and 100 kbp downstream of the PAM at the *B2M* and *TRAC* loci. For off-target analysis in the CRISPR-Cas3 system, probes were designed to cover 9 kbp upstream and 1 kbp downstream of each POT site.

Genomic DNA was extracted from Jurkat cells electroporated with Cas3-Cascade RNP complexes. DNA libraries were prepared using the SureSelect XT reagents and custom probe sets (Agilent Technologies). High-throughput sequencing was performed using the HiSeq X system (2 × 150 bp paired-end reads; Macrogen Japan). Raw sequencing reads were mapped to the GRCh38/hg38 reference genome using the BWA-MEM algorithm.

For on-target analysis, split reads and discordant read pairs were extracted using LUMPY [30] and SAMtools [31], respectively. The number of split reads per region was quantified with BEDtools [32]. Genomic loci where the ratio of split read counts (Cas3-treated sample vs. control) exceeded 5 were defined as cut points. Deletion patterns were then inferred based on discordant read pairs at each cut point. Off-target effects were analyzed as described previously [7]. Briefly, split read counts at each POT site in cells transfected with Cas3-Cascade RNP without crRNA were subtracted from those in cells transfected with the test crRNAs. For CRISPR-Cas9 off-target analysis, data were processed using CRISPResso2 [19].

### Statistical Analysis

All statistical analyses were performed using GraphPad Prism software (GraphPad Software, San Diego, CA). Specific statistical tests used are indicated in the figure legends. A P-value of less than 0.05 was considered statistically significant.

## Data Availability

All data produced in the present study are available upon reasonable request to the authors.

## Limitations of the study

Due to the lack of appropriate animal models for evaluating immune responses to human cells, all experiments in this study were conducted in vitro. Therefore, it remains to be determined in clinical trials whether Cas3-mediated knockout of *B2M* and *TRAC* will effectively reduce graft-versus-host responses and immune rejection in vivo. Nevertheless, accumulating evidence supports the notion that the CRISPR-Cas3 system may offer safety advantages over the Cas9 system. Our findings provide a solid foundation for further advancing the development of allogeneic T cell therapies using CRISPR-Cas3.

## Acknowledgments

We acknowledge the IMSUT FACS Core Laboratory for flow cytometry analysis and Rena Yamauchi for technical assistance. This research was supported from AMED (JP19am0401022 and JP23bm1223009h0001) and JSPS KAKENHI (23H00367).

## Author contributions

T.F., K. Yokoyama, and S. Watanabe conducted and analyzed most of the experiments and wrote the manuscript. Y.S. performed immunological analyses and contributed to manuscript writing. K. Yoshimi provided scientific discussion. K. Takeshita prepared the CRISPR-Cas3 proteins. K. Tamada and T.M. conceived and supervised the study and wrote the manuscript.

## Supplementary Figure legends

**Figure S1. Schematic overview of the experimental procedure for genome editing in human primary T cells.** PBMCs were activated with anti-CD3 antibody (OKT3), Retronectin, and IL-2 for three days, followed by electroporation with Cas3-Cascade mRNAs and crRNAs. Disruption efficiency, cell viability, and live cell number were analyzed on day 7.

**Figure S2. Off-target analysis in human primary T cells introduced with the CRISPR-Cas9 mRNA.** (A) Cells edited with Cas9_TRAC sgRNA. (B) Cells edited with Cas9_B2M sgRNA. (C, D) IGV images showing edited (top) and control (bottom) sequencing reads at off-target sites identified in *TRAC*-edited (C) or *B2M*-edited (D) human primary T cells. Detected off-target loci include the *ECT2L* gene locus on chromosome 6 and a locus on chromosome 14, highlighting the potential for unintended editing by CRISPR-Cas9.

