## Supplemental Figures 1-2 for "Efficient gene disruption with CRISPR-Cas3 in human T cells"

### Slide 1
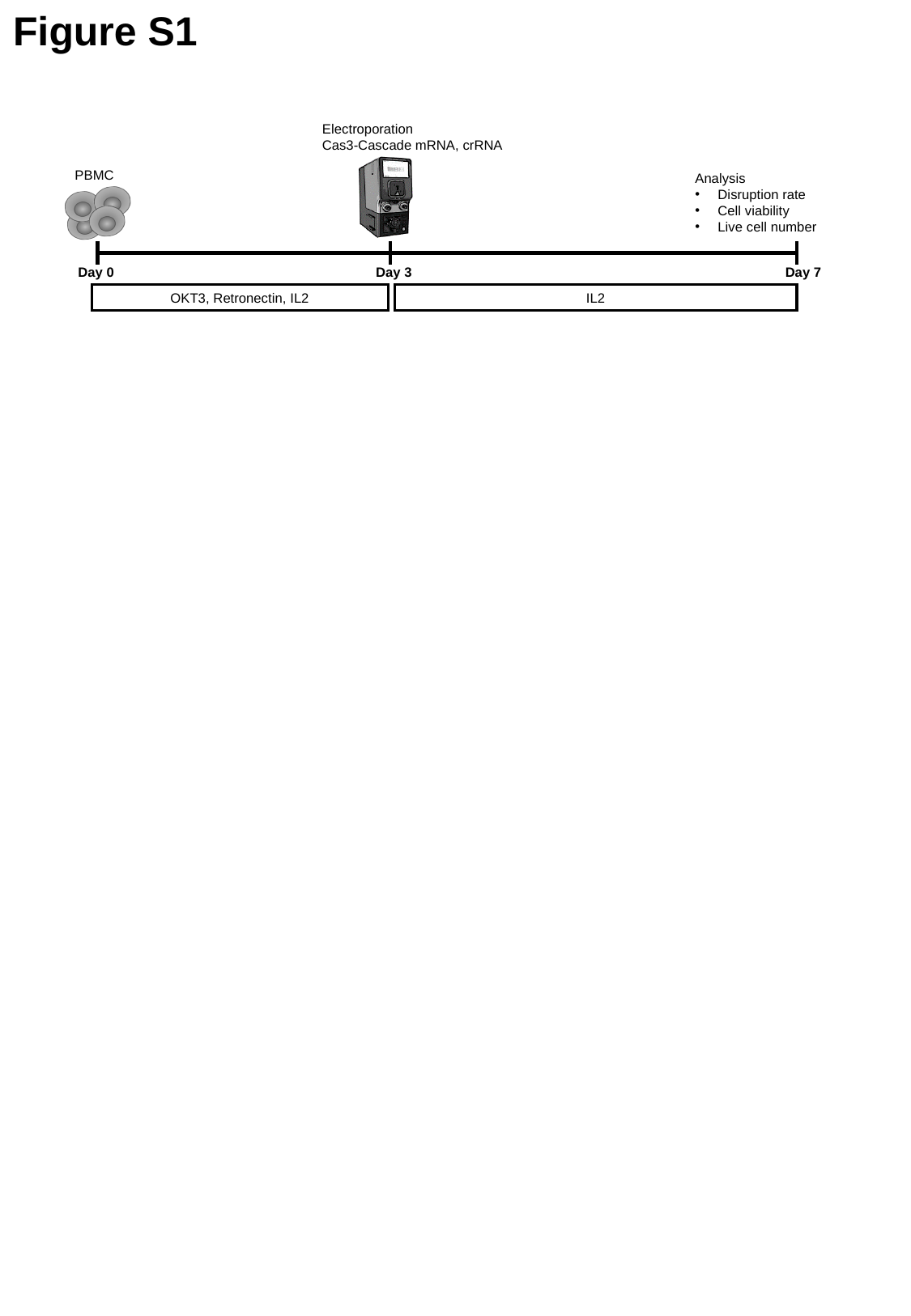

Figure S1
Electroporation
Cas3-Cascade mRNA, crRNA
PBMC
Analysis
Disruption rate
Cell viability
Live cell number
Day 0
Day 3
Day 7
OKT3, Retronectin, IL2
IL2

### Slide 2
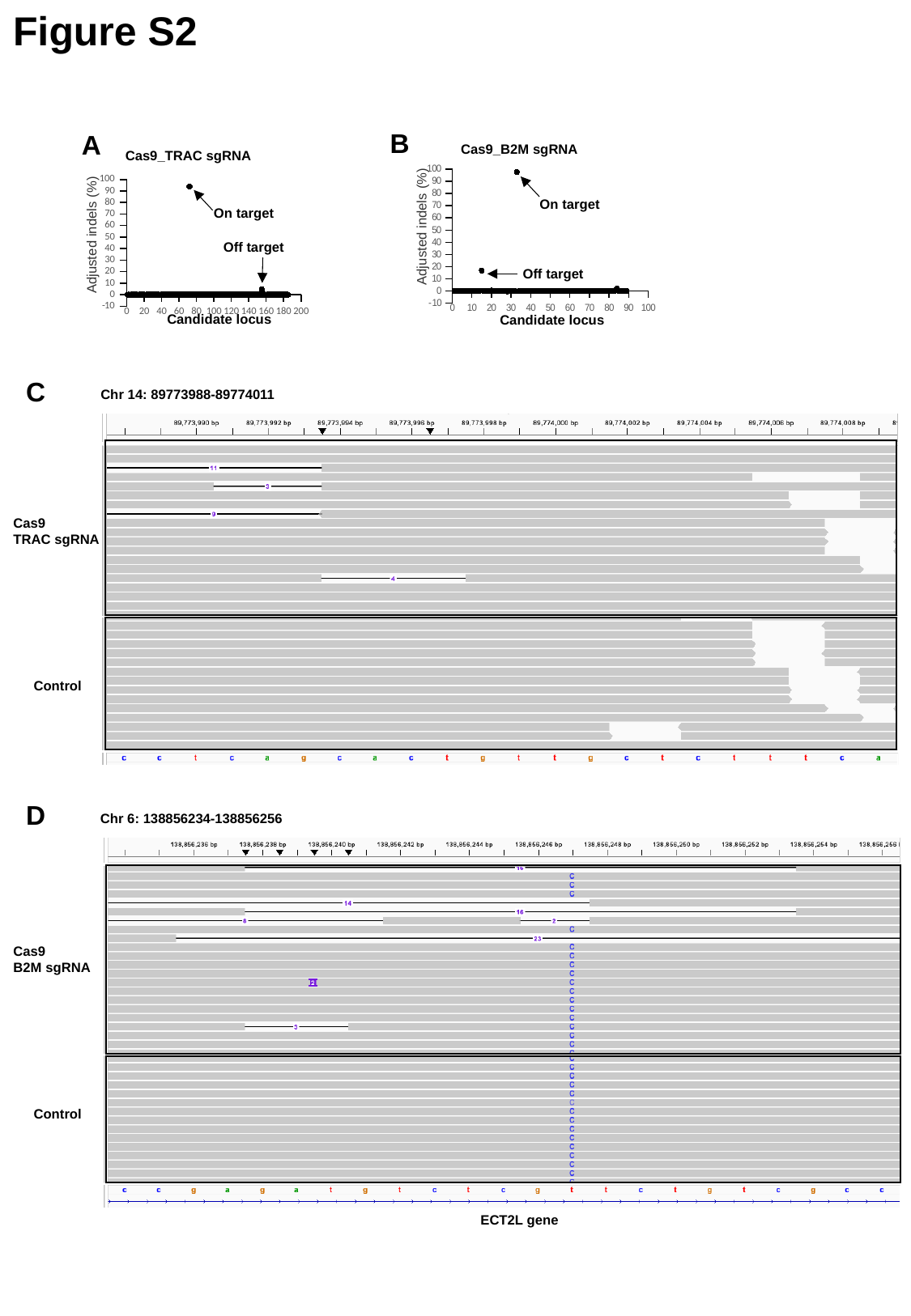

Figure S2
B
A
Cas9_B2M sgRNA
Cas9_TRAC sgRNA
#### Chart
| Category |
|---|
#### Chart
| Category | |
|---|---|On target
On target
 Adjusted indels (%)
 Adjusted indels (%)
Off target
Off target
Candidate locus
Candidate locus
C
Chr 14: 89773988-89774011
Cas9
TRAC sgRNA
Control
D
Chr 6: 138856234-138856256
Cas9
B2M sgRNA
Control
ECT2L gene
